# Improved diagnostic prediction of the pathogenicity of bloodstream isolates of *Staphylococcus epidermidis*

**DOI:** 10.1101/2020.10.16.342238

**Authors:** Shannon M. VanAken, Duane Newton, J. Scott VanEpps

## Abstract

With an estimated 440,000 active cases occurring each year, medical device associated infections pose a significant burden on the US healthcare system, costing about $9.8 billion in 2013. *Staphylococcus epidermidis* is the most common cause of these device-associated infections, which typically involve isolates that are multi-drug resistant and possess multiple virulence factors. *S. epidermidis* is also frequently a benign contaminant of otherwise sterile blood cultures. Therefore, tests that distinguish pathogenic from non-pathogenic isolates would improve the accuracy of diagnosis and prevent overuse/misuse of antibiotics. Attempts to use multi-locus sequence typing (MLST) with machine learning for this purpose had poor accuracy (~73%). In this study we sought to improve the diagnostic accuracy of predicting pathogenicity by focusing on phenotypic markers (*i.e*., antibiotic resistance, growth fitness in human plasma, and biofilm forming capacity) and the presence of specific virulence genes (*i.e., mecA, ses1*, and *sdrF*). Commensal isolates from healthy individuals (n=23), blood culture contaminants (n=21), and pathogenic isolates considered true bacteremia (n=54) were used. Multiple machine learning approaches were applied to characterize strains as pathogenic *vs* non-pathogenic. The combination of phenotypic markers and virulence genes improved the diagnostic accuracy to 82.4% (sensitivity: 84.9% and specificity: 80.9%). Oxacillin resistance was the most important variable followed by growth rate in plasma. This work shows promise for the addition of phenotypic testing in clinical diagnostic applications.

## INTRODUCTION

Coagulase-negative staphylococci (CoNS) are the most commonly isolated bacteria in clinical microbiology laboratories(1); however, they are not routinely considered pathogenic. CoNS, particularly *Staphylococcus epidermidis*, are among the most numerous commensal bacteria on the skin and mucous membranes(2). Although considered less pathogenic, CoNS are the most common opportunistic implanted medical device colonizers(3–6). Studies have shown that approximately 20% of all healthcare associated infections are caused by CoNS, where 90% of those are primary bloodstream infections(7). *S. epidermidis* also frequently contaminates otherwise sterile blood cultures during the blood draw(8, 9). As such, growth from blood cultures may indicate a false-positive resulting from skin contamination or a clinically significant bloodstream infection(8).

Published studies on the criteria for differentiating true from contaminant isolates of CoNS are conflicting, with many reporting a mixture of additional molecular typing methods required to improved diagnostic power(10–12). The Centers for Disease Prevention and Control clinical criteria for true positives require the detection of CoNS from two or more blood cultures within 48 hours and accompanying symptoms in the patient (*i.e*., fever, chills, and/or hypotension). However, these criteria alone are only acceptable as a screening tool. Laboratory confirmed bloodstream infection criteria and clinical signs combined are still insufficient to further distinguish bacteremia from contamination(13). Rapid multiplex PCR on suspected pathogenic isolates reduced unnecessary antibiotic use by identifying likely contaminating species, however it offers no way to distinguish pathogenic from commensal *S. epidermidis* isolates(14). The development of a superior discriminatory tool to distinguish pathogenic from non-pathogenic blood culture isolates of *S. epidermidis* would help prevent overuse and/or misuse of antibiotics, which contributes to the ongoing threat of antibiotic resistance. We begin by reviewing potential mechanisms for evaluating the pathogenicity of *S. epidermidis*.

*S. epidermidis* isolated from infections have been suggested to be a molecular subset of those found on the skin surface (*i.e*., they continue to carry out original functions of the non-infectious ‘lifestyle’) (15–17). This implies that, rather than passive infection, there may be certain lineages or specific virulence factors associated with the emergence of pathogens from a background of harmless ancestors. Specifically, conferred pathogenicity is manifested in remaining competitive through selection, namely antibiotic treatment, host microenvironment, or immune clearance(18). Virulence genes, including antibiotic resistance, biofilm formation capacity and other metabolic advantages, are commonly passed on pathogenicity island plasmids exogenously or through horizontal gene transfer. Genotype testing has been used to help identify isolates that carry specific virulence genes(19), and thus have potential to express virulence. Genotyping, however, does not take into consideration the gross gene pool and how many of those individual genes are redundant(20); nor does it account for variations in gene expression, post translational processing, or emergence of new gene mutations that affect ultimate phenotype. Therefore, phenotypic measures may serve as a better diagnostic tool(21, 22).

Biofilm formation is an exceedingly important aspect of staphylococcal growth and pathogenicity(23). *Staphylococcus* adheres strongly to host proteins in the skin, allowing it to live as a commensal bacterium. During surgical procedures, bacteria use surface adhesion proteins to adhere to deep tissues and onto implanted medical devices(24). These attributes contribute significantly to opportunistic infections and allow specific sub-populations the ability to invade the bloodstream(25). Genes conferring biofilm formation capacity have been suggested to be present in pathogenic and not in commensal CoNS isolates(5, 26). Examples include, SdrF, a surface protein that mediates attachment to keratin and collagen(27), and SesI, another cell-wall-anchored protein with as of yet undefined binding target(28). However, it is clear that targeting individual genes based upon product amplification methods would be ineffective, as the sources of disease recombine and are found in multiple genetic backgrounds(29).

Studies cataloguing virulence factors from clinical isolates are abundant. *MecA* is the most common virulence factor reported, especially in reference to methicillin-resistant *Staphylococcus aureus* (MRSA)(30–32). Encoding an alternative penicillin-binding protein, *mecA* confers resistance to a large class of antibiotics. It is a crucial part of the SCCmec cassette, the most widely disseminated staphylococcal mobile genetic element(31, 33, 34). SCCmec cassettes are found mostly in *S. aureus* and carry mix-and-match functional and virulence genes; however they have also recently been found in pathogenic and commensal CoNS isolates(35). Antibiotic overuse and/or misuse leads to further selection of these pathogenicity islands(36, 37). A resistance or virulence factor profile may be a possible diagnostic tool as presence of specific SCCmec cassettes may confer definitive patterns in resistance as well as other virulence genes and resulting phenotypes.

In addition to the suggested virulence factors, fundamental metabolic processes of bacteria are also recognized as a prerequisite for disease(38). Microbial fitness during pathogenesis requires efficient utilization of available nutrients. Strategies for overcoming the nutrient-limited environment *in vivo* include, upregulation of peptide or amino acid transport mechanisms and proteins that enable the acquisition of nutrients sequestered by the host(38, 39). Considering CoNS easily contaminate blood cultures, examining growth rates of isolates in human blood fractions may provide invaluable differentiation between commensal and pathogenic strains(38).

One molecular strategy that has been used to interpret *S. epidermidis* identified in blood culture is multi-locus sequence typing (MLST)(8, 40, 41). In one study, machine learning algorithms were employed by treating the MLST as a 7-dimensional variable-space(41). The genetic variability was substantial between clinical *S. epidermidis* isolates, with 44 different sequence types found in 100 isolates(41). However, the best machine learning model only yielded 73% diagnostic accuracy. The overlap between clinically significant isolates and those representing sample contamination appeared to prevent the exclusive use of MLST in the clarification of blood cultures recovering *S. epidermidis*(41).

The ongoing investigation and continued controversy in distinguishing pathogenic from non-pathogenic isolates of *S. epidermidis* indicate that no single feature, factor, or test is sufficient for this task. To that end, we sought to develop a panel of predominantly phenotypic tests that can rapidly be deployed in standard clinical microbiology laboratories for the purpose of diagnosing clinically significant *S. epidermidis* bacteremia. On a library of isolates defined as either commensal, contaminant, or pathogenic we evaluated the potential for antibiotic susceptibility, biofilm formation capacity, growth fitness in plasma, and some select virulence genes to accurately differentiate these isolates. Machine learning algorithms were applied to the panel in aggregate to predict the pathogenicity and determine the most important predictive features.

## METHODS

### Clinical isolate collection

All isolates were previously obtained for a prior study(41)and were graciously provided by Dr. John G. Younger. Specifically, commensal isolates (n=23) were obtained from healthy volunteer thumbprints on mannitol salt agar plates. Pathogenic isolates (n=54) were recovered by the Michigan Medicine Clinical Microbiology Laboratory from positive blood cultures detected using the BacT/Alert/FAN system (bioMérieiux, Durham, NC). Each of these isolates were present in two sets of blood cultures obtained contemporaneously in symptomatic patients indicating “true-positives”. Isolates classified as contaminants (n= 21) were those that were detected in only one set of culture bottles and therefore considered “false-positives”. Cultures for all experiments were only 1-2 culture passages past original isolation state to reduce elimination of elements not required for standard growth.

### Antibiotic susceptibility

Susceptibility testing was performed *via* disk-diffusion on Mueller Hinton Agar with antibiotic disks obtained from Oxoid (ThermoFisher, Lenexa, KS). Standard criteria for interpretation of results were applied as outlined by Clinical Laboratory Standards Institute (CLSI)(42).

### Biofilm formation

Biofilm formation was evaluated by crystal violet staining after 24 hours of biofilm growth in 96-well plates. Briefly, single colonies were grown overnight in tryptic soy broth supplemented with 1% glucose (TSBg) at 37°C. Cultures were then diluted to an optical density at 600 nm (OD_600_) of 0.5-0.6 followed by a 1:100 dilution to inoculate ~5×10^5^ cells per well in triplicate. Biofilms were grown at 37°C for 24 hours and then rinsed to remove nonadherent cells. Biofilms were stained with 0.2% crystal violet (CV) for 15 minutes then rinsed 3 times in tap water. Stained biofilms were dried on the benchtop for 16-20 hours and then incubated at room temperature with 33% acetic acid for 15 min. Samples were moved to a clean plate then absorbance was measured at 570 nm (A_570_). All values were normalized to the strong biofilm former *S. epidermidis* RP62a (ATCC 35984) to account for significant day-to-day variation.

### Growth fitness

Growth curves were obtained by time course measurement of OD_600_ in TSBg alone and TSBg supplemented with 10% human plasma. Single colonies were grown overnight in TSBg at 37°C then diluted to OD_600_ of 0.5-0.6 and a 1:100 dilution was used to achieve initial inoculum of 5×10^5^ cells per well. Growth curves for each isolate in each medium were obtained in at least triplicate. Curves were fit to a Gompertz function to determine the maximum growth rate (μ), lag phase (λ), and maximum OD_600_ (A) (see **Supplemental Figure S1** and **Supplemental Table S1**). The ratio of each parameter (plasma:no plasma) are reported.

### Virulence genes

The presence of virulence genes was determined by PCR on 5 ng of genomic DNA and was considered positive or negative based on presence or absence of a single product band per reaction; observed on 1% agarose gel in TAE buffer. Genomic DNA was isolated from overnight cultures using Promega Wizard Genomic DNA Isolation kit (Madison, WI), according to the manufacturer’s instructions for *Staphylococcus spp*. Primer sequences are listed in Supplemental Table S2.

### Statistical analysis

For antibiotic resistance data we determined the frequency of resistance for each antibiotic grouped by isolate type (commensal, contaminant, and pathogen). To create a binary phenotype, intermediate resistance, determined from the zone of inhibition assay, was treated as resistant. Hypothesis testing for antibiotic resistance was performed using the χ^2^ test where the null hypothesis was that each isolate type had the same frequency of resistance. Significance was based on a p-value of less than 0.05 with a Bonferonni correction for multiple comparisons.

For biofilm formation capacity, comparisons were made by Kruskall-Wallis analysis of variance followed by post-hoc pair-wise Dunn’s test for comparisons due to the nonparametric nature of the data.

For growth fitness we were interested in differential growth parameters in the presence and absence of plasma as a surrogate for growth fitness in blood. These differential parameters were presented as ratios of the raw growth parameters (see methods and supplementary materials) with and without plasma (plasma:no plasma). To demonstrate the overall variance in data, the ratios for each isolate, grouped by type, are shown and then summarized by type. Shapiro tests for normality were performed on each parameter ratio. For those that were assumed to be normally distributed, ANOVA with post-hoc pairwise Tukey tests were performed. For those that could not be assumed to be normally distributed, Kruskall-Wallis with post-hoc Dunn’s test were performed.

For virulence genes, the frequency with which the gene is present within isolate types was determined and hypothesis testing using the χ^2^ test with Bonferroni correction for multiple comparisons was performed like that for the antibiotic resistance.

Given that we evaluated methicillin resistance from both a genotypic and phenotypic perspective, we chose to compare these methods by generating and comparing confusion tables for each isolate group.

### Predictive modeling

We initially performed an analysis of similarity (ANOSIM) which demonstrated that dissimilarity between isolates types was higher than that within each type but quite similar to that of the commensal and contaminates types (**Supplemental Figure S2**). The pathogen type however had significantly less dissimilarity indicating the possibility that this isolate type is a more homogenous subgroup. Therefore, we chose to proceed under the assumption that isolates characterized as commensal or contaminant should be considered to be from one population and the pathogens from another. Therefore, all predictive modeling henceforth will be for the binary pathogen *vs* non-pathogen phenotype with the commensal and contaminant types combined to form the non-pathogen phenotype. A total of five predictive models were developed: generalized linear model for binary logistic regression (glm), support vector machine (svm), recursive partitioning (rpart), conditional inference tree (ctree), and a random forest from conditional inference trees (cforest).

For the glm model, we performed a forward and backward stepwise selection of variables based on the Akaike Information Criterion (AIC). For the svm model, we used a radial basis function and tuned the gamma and cost for best performance. For the rpart model, the size of the tree was determined by choosing the complexity parameter cutoff that minimizes the cross-validation error. The ctree splits were based on the raw value of the test statistic with a maximum depth of 3. The cforest model implements a random forest from the conditional inference trees above. To evaluate the predictive performance, the dataset was randomly divided such that 75% of the samples were used to train the model and 25% were used to test the model. Area under the receiver operator curves (AUROC) were determined as well as the accuracy, sensitivity, and specificity of each model. This was repeated five times for cross-validation and values are reported as mean and standard deviation. For each optimized model, the specific variables of importance were determined and the number of models in which they occur are presented.

## RESULTS

### Antibiotic susceptibility

*S. epidermidis* clinical isolates were tested for susceptibility to 9 commonly used antibiotics (Figure 1). In general, there was a high frequency of antibiotic resistance in the commensal isolates supporting the concept of *S. epidermidis* being a universal reservoir for antibiotic resistance genes(43). Resistance to ciprofloxacin, oxacillin, and doxycycline was significantly different between the isolate types suggesting that these traits may be useful in predicting isolate type.

**Figure 1:**
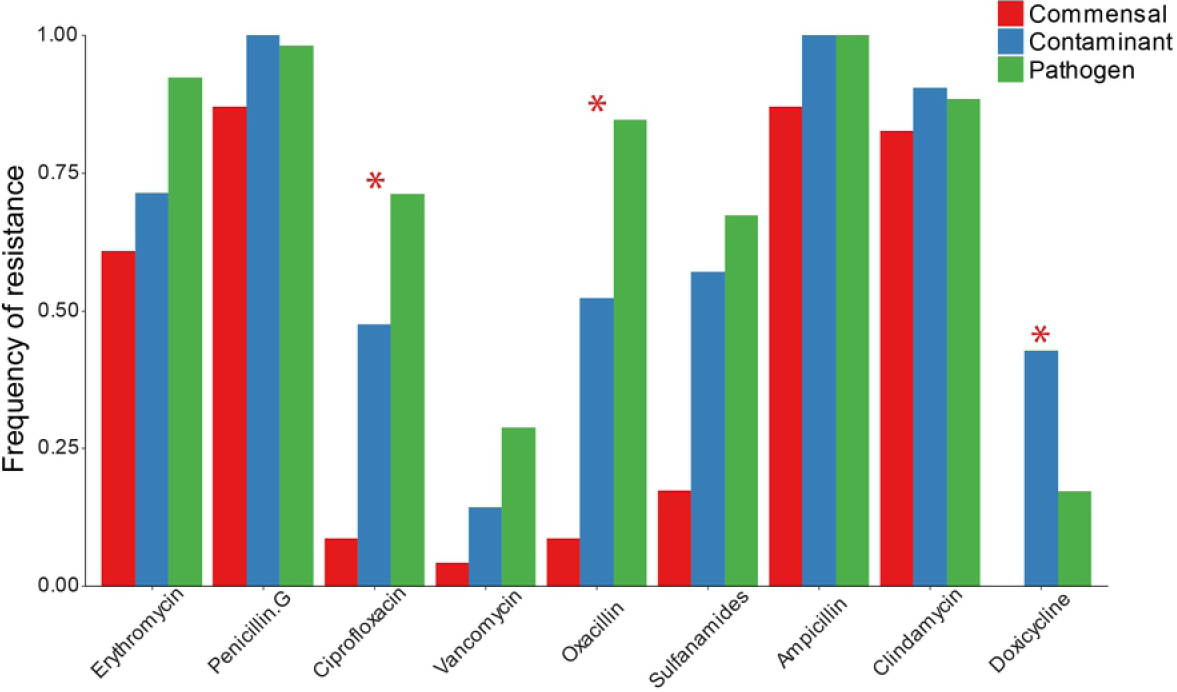
Frequency of Antibiotic resistance among commensal (n=23), contaminant (n=21), and pathogenic (n=54) *S. epidermidis* clinical isolates. * Indicates χ^2^ p<0.05 with Bonferroni correction for multiple comparisons.

### Biofilm formation

Biofilm formation is a commonly described virulence factor for *S. epidermidis*. There was high variability in biofilm forming capacity within isolate types. While the pathogen type had a greater median crystal violet staining, the difference was not statistically significant (Figure 2).

**Figure 2:**
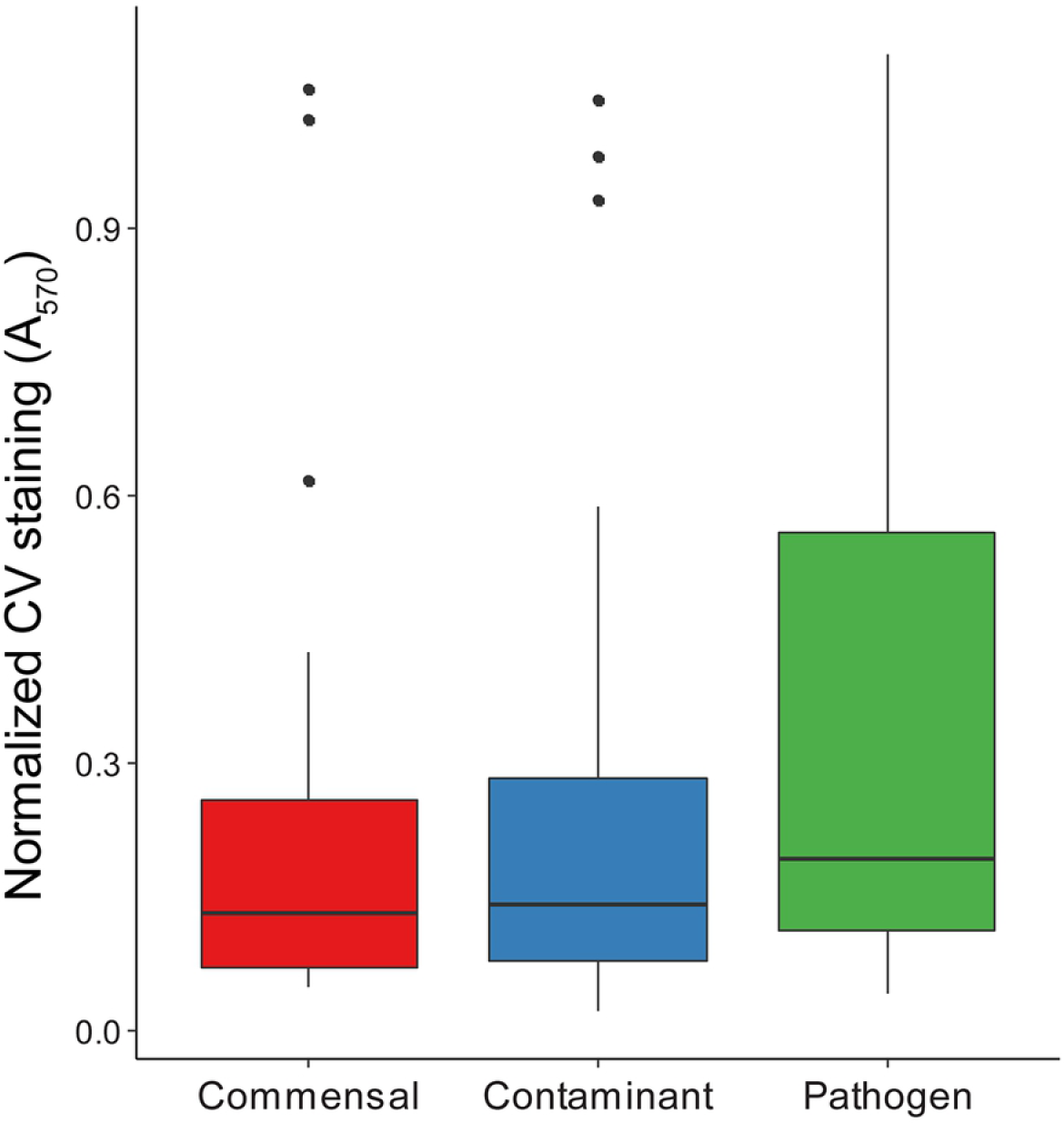
Biofilm forming capacity of isolates as determined by crystal violet staining normalized to the established biofilm former *S. epidermidis* RP62A.

### Growth fitness

We hypothesized that pathogenic isolates grow better in human plasma when compared to non-pathogenic isolates. This is based on the idea that transition to the new environment of the bloodstream favors pathogenic strains over commensals. In most cases, a delay in lag time (λ) was seen in all isolates grown in plasma, suggesting overall inhibition. However, commensal isolate growth seemed to be most negatively affected by the presence of plasma. This was manifested in increased lag time (λ) and decreased growth rate (μ) for the commensal isolates relative to the contaminants and pathogens (Figure 3). Pathogenic isolates tended to be less inhibited in plasma compared to both contaminants and skin isolates, suggesting an increased fitness in blood. While on average all three groups grew faster without plasma, skin isolates had the greatest average inhibition in plasma. ANOVA for differences between types for growth rate (μ) ratio had p<0.05 but all post-hoc pairwise comparisons were not significant. On average, pathogens have the shortest lag time (λ) ratio. This suggests they have the greatest fitness in plasma during early growth. Commensal isolates are the most inhibited during early growth in plasma. Contaminants were similar in lag time (λ) ratio to pathogens, with a slightly higher average ratio, however the contaminant group was not statistically different than either commensals or pathogens. Kruskal-Wallis with post-hoc Dunn’s test has p<0.05 for commensal *vs* pathogen.

**Figure 3:**
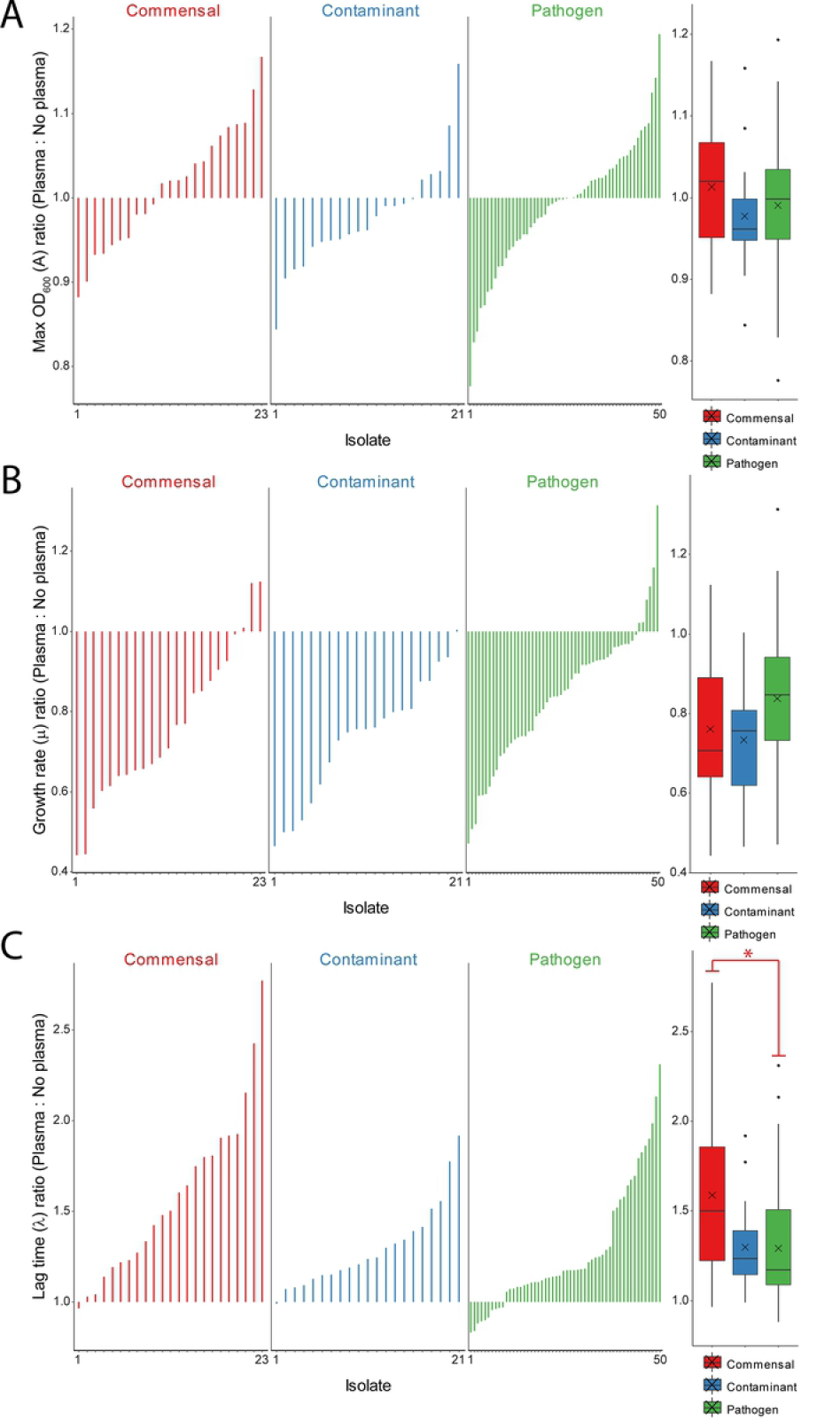
For each isolate the ratio (plasma:no plasma) for the **(A)** maximum OD_600_ (A), **(B)** growth rate (μ) and (C) lag time (λ) are plotted. To the right of each plot the data is summarized a box and whisker plot that includes the mean denoted with an X. There were no statistical differences for the maximum OD_600_ (A) ratio. For the maximum growth rate (μ) in **(B)** the data was normally distributed and an ANOVA had a p<0.05 but there were no significant post-hoc pairwise differences. For the lag time (λ) ratios in **(C)** the data could not be assumed as normally distributed. Kruskall-wallis with post-hoc Dunn’s test had p<0.05 when comparing the commensals to the pathogens (denoted by *).

### Virulence genes

*MecA* was present in all types, with the largest frequency of 94.4% in the pathogenic isolates. Contaminants had an intermediate *mecA* frequency (80%), which was significantly higher than the frequency for skin isolates (34.8%). An increase in frequency from commensal to contaminant to pathogen was seen for *sdrF*. Only mecA was statistically significant, though (Figure 4).

**Figure 4:**
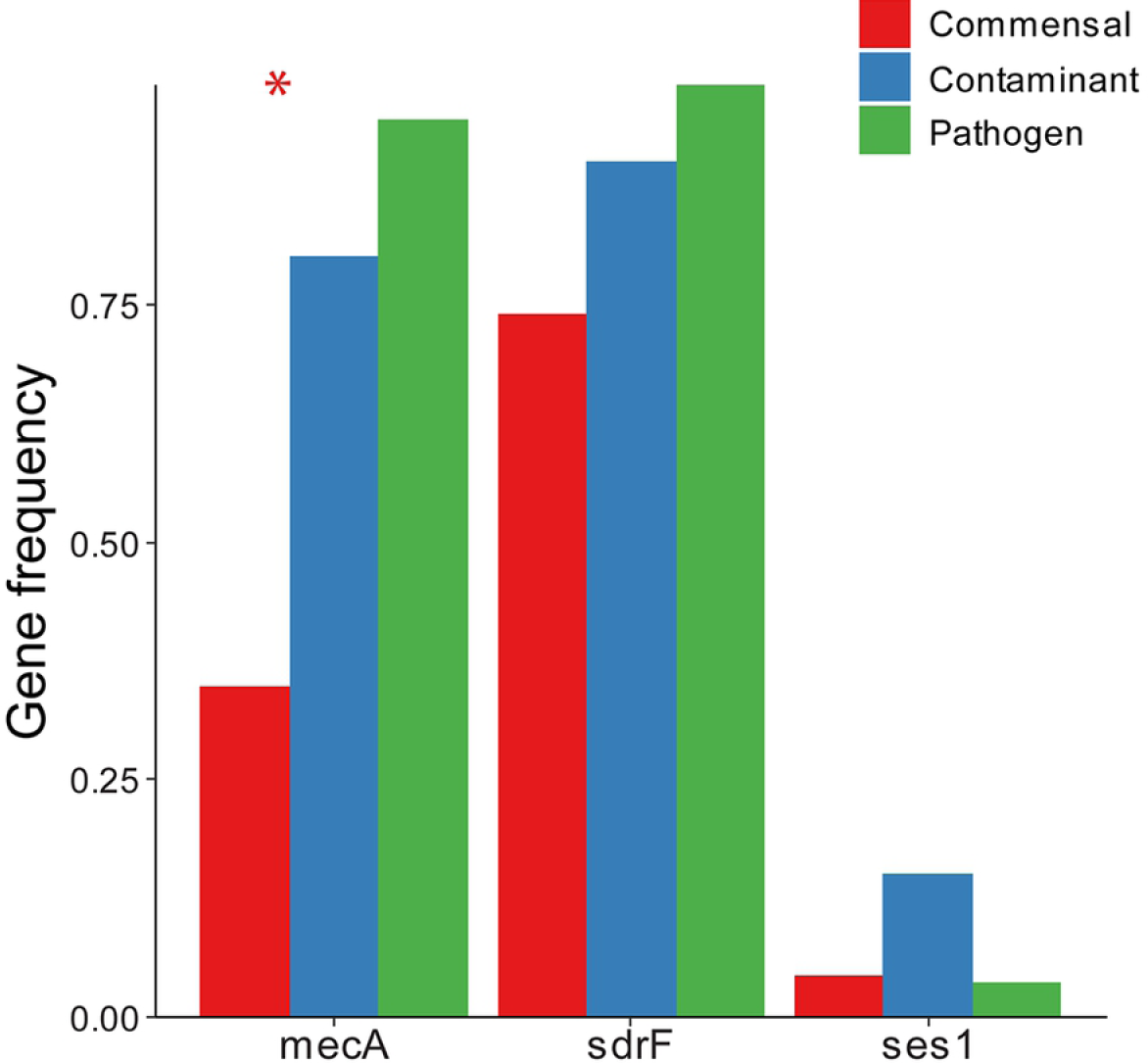
Frequency of virulence genes. * indicates χ^2^ p<0.05 with Bonferroni correction for multiple comparisons.

As noted earlier, there is the potential for discordance between genotypic and phenotypic traits. This is most illustrative, clinically, when considering the presence of the *mecA* gene as a surrogate for methicillin resistance. Certainly if the *mecA* gene is detected, although not phenotypically expressed initially, it could be induced with exposure to oxacillin. However, if the gene is not detected the isolate could still have phenotypic resistance. This “false negative” could result in under treatment of a critically ill patient. Since both measures were considered for each isolate in this study, we evaluated their discordance *via* 2×2 confusion tables (**Supplemental Figure S3**). The *mecA* gene was detected in 18 out of 71 isolates yet the isolate was phenotypically susceptible to oxacillin. However, there was only 1 isolate out of 23 where *mecA* was not detected but was resistant to oxacillin. When considering each isolate group separately, 8, 5, and 6 of the discordant events were in the commensal, contaminant, and pathogen groups respectively.

### Predictive modeling

Five predictive modeling algorithms were applied to the panel of tests performed here with the goal of differentiating pathogenic from non-pathogenic isolates. The performance was characterized by area under the receiver operator curves and calculated accuracy, sensitivity, and specificity (Table 1). All models with the exception of the generalized linear model (glm) outperformed the previous attempts that focused on MLST alone. For this dataset, the condition inference tree (ctree) model had the best performance. Pathogens were able to be differentiated from non-pathogens with 82.4% accuracy, 84.9% sensitivity, and 80.9% specificity. For future refinement and validation of these models it is important to consider the relative importance of each variable in the model. The frequency with which a particular variable appears in each validation of the five models is indicative of its predictive power and is shown in Figure 5. Oxacillin, vancomycin, and erythromycin phenotypic resistance and the growth rate (μ) ratio were the most frequent features in the models. Oxacillin resistance was present in all but one model.

**Table 1.** Predictive model performance

| Model | Accuracy | Sensitivity | Specificity |
| --- | --- | --- | --- |
| *glm* | 72.1 ± 11.6 | 76.2 ± 17.7 | 68.1 ± 29.2 |
| *svm* | 76.5 ± 1.8 | 73.7 ± 15.7 | 81.7 ± 11.9 |
| *rpart* | 76.7 ± 6.2 | 75.8 ± 16.8 | 79.6 ± 7.4 |
| *ctree* | **82.4 ± 4.2** | **84.9 ± 12.0** | **80.9 ± 6.7** |
| *cforest* | 78.7 ± 4.1 | 84.2 ± 16.8 | 77.1 ± 8.5 |

**Figure 5:**
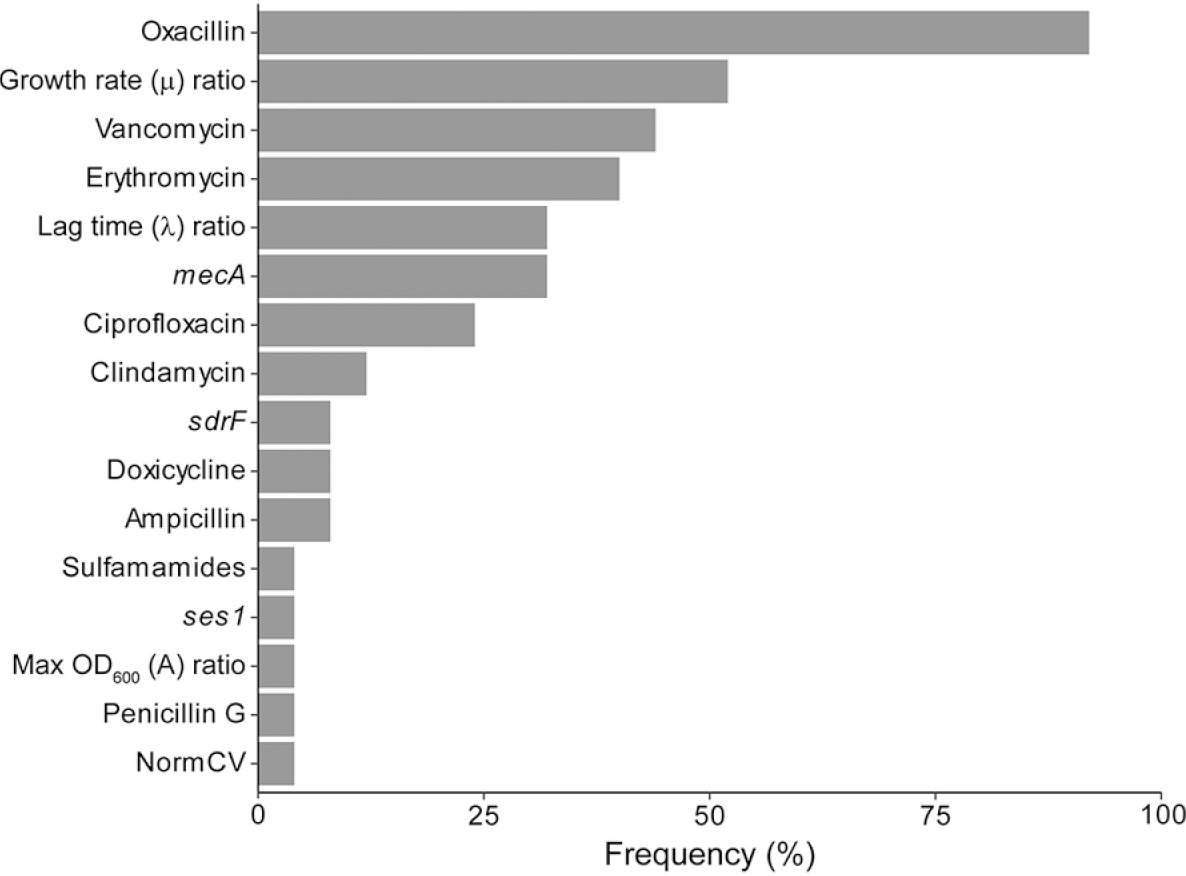
Frequency of specific variable appearance

## DISCUSSION

In this study, we compared three types of *S. epidermidis clinical* isolates (*i.e*., commensal, blood culture contaminants, and true pathogens) with respect to a panel of predominantly phenotypic tests. Our goal was to address an unmet need for more accurate diagnosis of clinically significant bloodstream infections caused by *S. epidermidis*. We begin with a summary of individual test results and their interpretations, followed by a discussion of the performance of the predictive models, and conclude with a discussion of the implications, limitations, and future directions for this work.

Pathogenic *S. epidermidis* isolates originate from patients in a healthcare setting where antibiotic treatment leads to potential selection of antibiotic resistance cassettes and pathogenicity islands(3). This phenomenon is highlighted in this study as pathogens and contaminants from hospitalized patients had consistently higher frequency of antibiotic resistance than commensal isolates from healthy individuals. In particular, pathogens were significantly more likely to be resistant to ciprofloxacin and oxacillin. In addition, commensal isolates had both reduced growth rates and increased time to exponential growth in the presence of blood plasma compared to contaminants and pathogens, as has been demonstrated previously(44). This could result from the fact that the commensal isolates were obtained from thumb prints of healthy patients rather than blood cultures of patients with suspected bloodstream infection and were never exposed to the blood microenvironment. There was significant variability in biofilm forming capacity within isolates types and no detectable difference between groups. All types of *S. epidermidis* isolates were able to build biofilms as is likely required for colonization of the skin(16). Pathogens, however, may have more overlapping biofilm formation mechanisms as part of the selection for pathogenicity islands.

While several of the individual tests performed demonstrated differential trends between groups, no one factor, genotypic or phenotypic, distinguished isolates with perfect accuracy. This is not entirely surprising as frequent horizontal gene transfers increase competition between selected clones carrying competing beneficial mutations by moving multiple selected sites into a common background(29). In addition, MLST variants evolve and new variants emerge over time indicating that MLST cannot be used reliably on its own. Given the already described drawbacks of genotypic testing, we incorporated phenotypic tests in addition to amplifying a few virulence genes known to differentiate pathogens from non-pathogens. Our best performing model (*i.e*., conditional inference tree) distinguishes non-pathogens from pathogens with 82.4% accuracy. This represents a significant improvement over the MLST model which had an accuracy of only 73%(41).

Overall review of all of the model results, indicates that the greatest determinants of pathogenicity were phenotypic methicillin, vancomycin, and erythromycin resistance and growth rate (μ) ratio (plasma:no plasma). Oxacillin resistance was the most significant determinate between the two isolate groups. Modeling using oxacillin resistance or *mecA* presence alone gave 80% accuracy with 91% specificity, consistent with previous findings(45). Interestingly, a previous study on k-mer mapping to the *mecA* gene was the best predictor for *S. epidermidis* isolates from infection, giving a classification accuracy of 75% on its own, also consistent with our findings(29). As has been shown with other studies(46), some isolates were mismatched when comparing genotypic with phenotypic methicillin resistance. A smaller percentage of pathogens (13%) than non-pathogens (30%) had discordant phenotypic *vs* genotypic methicillin resistance status. (recall **Supplemental Figure S3**). All but one of the discordances were seen in phenotypically susceptible isolates. This suggests that pathogens are more likely to develop phenotypic methicillin resistance through selection for and expression of a genetic element carrying the *mecA* gene. Absences of antibiotic treatment induced selection and infrequent healthcare exposure can explain less congruency seen in commensal isolates taken from healthy volunteers. These together suggest that phenotypic testing is exceedingly important as *mecA* presence alone is a poor indicator of phenotypic methicillin resistance especially in commensal isolates. Misdiagnosis of commensals as pathogenic strains contributes to inappropriate antibiotic treatment(47).

Contaminants are described as being a commensal strain introduced into the bottle because of poor skin decontamination during collection, suggesting that they are just a genetic subset of commensals. Based on our findings, this seems to be mostly correct. In almost all traits tested, contaminants consistently had an intermediate frequency of factors or results. Alternatively, the conditions in the culture bottle which include host blood may induce some characteristics of a pathogen in these ‘accidental’ contaminants. The virulence trait profile of each isolate seems to be on a spectrum from completely inert commensals to pathogens loaded with a large collection of virulence factors, with most contaminants lying in between. This highlights the need for complex machine algorithms to distinguish true pathogens from commensal isolates due to the highly complex array of virulence factors that confer pathogenicity. Determining criteria to correctly identify pathogens may be easier due to the general homogeneity of pathogenic isolates, while inconsistencies and variations in commensal and contaminant virulence traits make false positives inevitable.

*S. epidermidis* has been described as a reservoir of resistance and virulence determinants that can be mobilized into the microbial community(16). Study of this reservoir could provide an early warning system for future clinically relevant antibiotic resistance mechanisms or evolving virulence. Varying collections of resistance and pathogenic genes in each bacterium make genetic evidence challenging to interpret in a clinical situation. Observation of phenotypic expression of pathogenic traits, however, may help narrow down the search; rather than assessing the isolates’ potential for expression, it is testing if the virulence factor is being expressed(33, 48). Ultimately, evidence of increased presence of phenotypic virulence or resistance will improve the accuracy of diagnostic tools in the clinical setting and thereby provide a means to prevent further selection and dissemination through the microbial and human population(49).

Of note, some relationships observed in this study could be overrepresented due to lowered isolate counts in commensal and contaminant groups as compared to pathogens. By combining the commensal and contaminant groups for predictive modeling we were able to generate a balanced dataset. However, the entire sample was relatively small in general. An additional limitation for this study was the lack of clinical information related to each patient, including sex, age, diagnosis, previous or current antibiotic treatment, comorbidities, or presence of indwelling medical device. Such information is critically important in determining the risk for bloodstream infection(50, 51). Regional differences in community health complicate comparing results to similar studies from different areas. Inconsistencies between published virulence surveys are abundant(52–56)and can be attributed to variances in geographical location where local healthcare and population affect selection factors that impact virulence factor carriage.

Future validation testing is important to continue to link pathogenicity with both patient and bacterial factors that are easily determined. A multicenter trial could look for patterns of risk factors related to pathogenicity of CoNS healthcare associated infections. Patient health information, along with phenotypic and genotypic factors expressed by the *S. epidermidis* isolate, could compound into a library of data that machine learning algorithms can be trained on to distinguish when a clinically isolated *S. epidermidis* strain is a pathogen or not. This can affect each patient’s treatment in clinical practice, as discovering a pathogenic *S. epidermidis* isolate may provide insight into the general pathogenic traits possessed by each individuals’ skin microbiome. This level of targeted treatment could reduce the overuse and misuse of antibiotics as antibiotic resistance has emerged as one of the major urgent threats to public health(57). Antibiotic treatment has potential adverse outcomes with adverse drug reactions and hypersensitivity reactions accounting for more than 3% of hospital admissions(58), which generates a significant burden on the health care system through secondary effects(59).

## ACKNOWLEDGMENTS

This work was supported by National Institutes of Health (NIH; https://www.nih.gov/), National Institute of Allergy and Infectious Diseases (NIAID; https://www.niaid.nih.gov/), AI-128006 (JSV) as well as the A. Alfred Taubman Medical Research Institute, https://www.taubmaninstitute.org/ (JSV). The funders had no role in study design, data collection and analysis, decision to publish, or preparation of the manuscript

## AUTHOR STATEMENT

SMV contributed to methodology, investigation, formal analysis, and writing. DN contributed resources, data curation, and writing. JSV contributed conceptualization, methodology, formal analysis, project administration, supervision, funding, and writing. We have no conflicts of interest.

